# SeqRepo: A system for managing local collections biological sequences

**DOI:** 10.1101/2020.09.16.299495

**Authors:** Reece K. Hart, Andreas Prlić

**Affiliations:** biocommons, San Francisco, CA; Invitae, Inc. San Francisco, CA

**Keywords:** bioinformatics, sequence analysis, hgvs, uta, biocommons, ga4gh, refget

## Abstract

**Motivation:** Access to biological sequence data, such as genome, transcript, or protein sequence, is at the core of many bioinformatics analysis workflows. The National Center for Biotechnology Information (NCBI), Ensembl, and other sequence database maintainers provide methods to access sequences through network connections. For many users, the convenience and currency of remotely managed data are compelling, and the network latency is non-consequential. However, for high-throughput and clinical applications, local sequence collections are essential for performance, stability, privacy, and reproducibility.

**Results:** Here we describe SeqRepo, a novel system for building a local, high-performance, non-redundant collection of biological sequences. SeqRepo enables clients to use primary database identifiers and several digests to identify sequences and sequence alises. SeqRepo provides a native Python interface and a REST interface, which can run locally and enables access from other programming languages. SeqRepo also provides an alternative REST interface based on the GA4GH refget protocol.

SeqRepo provides fast random access to sequence slices. We provide results that demonstrate that a local SeqRepo sequence collection yields significant performance benefits of up to 1300-fold over remote sequence collections. In our use case for a variant validation and normalization pipeline, SeqRepo improved throughput 50-fold relative to use with remote sequences. SeqRepo may be used with any species or sequence type. Regular snapshots of Human sequence collections are available.

It is often convenient or necessary to use a computed digest as a sequence identifier. For example, a digest-based identifier may be used to refer to proprietary reference genomes or segments of a graph genome, for which conventional identifiers will not be available. Here we also introduce a convention for the application of the SHA-512 hashing algorithm with Base64 encoding to generate URL-safe identifiers. This convention, *sha512t24u*, combines a fast digest mechanism with a space-efficient representation that can be used for any object. Our report includes an analysis of timing and collision probabilities for *sha512t24u*. SeqRepo enables clients to use sha512t24u as identifiers, thereby seamlessly integrating public and private sequence sets.

**Availability:** SeqRepo is released under the Apache License 2.0 and is available on github and PyPi. Docker images and database snapshots are also available. See https://github.com/biocommons/biocommons.seqrepo.

## Introduction

Many bioinformatics analysis pipelines require access to biological sequence data. One example is genetic variation data, which requires access to all sequences that are used as references in order to validate sequence bounds and to normalize variants (den Dunnen et al., 2016; Tan et al., 2015).

A typical whole genome sequencing sample has between 3.5 and 12 million variants (Hwang et al., 2019). Variant analysis pipelines for data of such volume need fast, random access to an assortment of genome, transcript, and protein sequences. While network-accessible databases (Ruffier et al., 2017; Sayers, 2010) are convenient, the latency is prohibitive for this high-throughput setting. Furthermore, dependencies on remote services create risks for privacy, reproducibility, and overall system availability. These were the problems for which we developed SeqRepo in 2016 as a component for the hgvs Python package (Wang et al., 2018). Using SeqRepo increases validation and variant projection throughput by nearly 50-fold relative to remote sequence access. We are unaware of tools similar to SeqRepo that enable the management and efficient distribution of sequence collections.

Sequence databases are often highly redundant within and between data providers. Deduplication of sequence datasets may be efficiently achieved using a digest or hash algorithm, such as SEGUID for protein sequences (Babnigg & Giometti, 2006). Here, we propose an approach that uses a variation of the SHA-512 hashing algorithm (National Institute of Standards and Technology, 2015) with Base64 encoding (Josefsson & Others, 2006) to generate URL-safe identifiers. The sha512t24u convention is a conceptual successor to SEGUID, but is computed approximately twice as fast on 64-bit processors and may be used in URLs without encoding. (SEGUID uses characters that are reserved for URL delimiters.) The GA4GH Variation Representation Specification (Babb et al., 2020) and the GA4GH refget protocol (GA4GH, 2019) have adopted the sha512t24u convention to generate computed object identifiers.

SeqRepo, presented here, provides these features:

1. Deduplication and compression of sequences;
2. Fast random access to sequences and sub-sequences;
3. Space-efficient storage and transfers of snapshots;
4. Ability to use of conventional, primary database identifiers and digest-based identifiers;
5. Publicly accessible snapshots of Human sequences;
6. Privacy and availability benefits appropriate for use in clinical settings.

## Implementation

### The sha512t24u Digest

We sought a digest and encoding that is based on existing standards, that may be implemented in prevalent programming languages, and may be readily used in URLs. Our method, dubbed *sha512t24u*, constructs a SHA-512 binary digest, truncated to 24 bytes (192 bits), and represented using “base64url” encoding. Figure 1 shows an implementation in Python; sha512t24u has also been implemented in Go, Java, javascript, and Perl.

**Figure 1.**
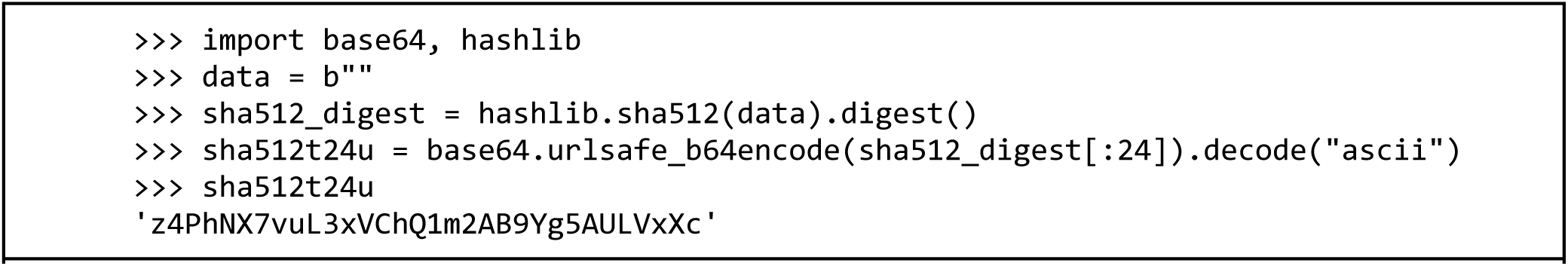
The sha512t24u digest in Python.

### SeqRepo Library

SeqRepo consists of two components: a module, fastadir, that stores sequences non-redundantly using an internal key, and a second module, seqaliasdb, that stores identifiers associated with the internal key. SeqRepo is written in Python 3.

#### Sequence Module: fastadir

Sequences are stored in FASTA-formatted files and index using Blocked GZipped Format (BGZF) (Li, 2011). Fast random access to BGZF files is provided by the PySAM library (PySAM Developers, n.d.).

When loading a sequence into SeqRepo, an internal *sequence identifier* is generated based on the *sha512t24u* convention. If the sequence does not already exist, it is written in a FASTA file with a timestamped filename using the internal sequence identifier as the FASTA identifier. A sequence manifest database, implemented in SQLite, stores sequence length, alphabet, and the path to the BGZF file that contains the sequence. Sequences in SeqRepo are immutable and not deletable; therefore, sequence files never change and exist in all future SeqRepo snapshots. New releases require only the space for new sequences and manifest db.

#### Alias Module: seqaliasdb

Once a new or existing sequence is confirmed in the sequence database, SeqRepo loads identifiers for the sequence as provided by the client. Identifiers are stored in a second SQLite database as <*namespace, alias*> pairs and are associated with the internal sequence identifier. Namespaces from <u>identifiers.org</u> are used when available. Aliases are required to be unique within a namespace. If an <*namespace, alias*> pair already exists, SeqRepo verifies that the associated sequence identifier matches the sequence being loaded; if it does not, the older association is deprecated in the database and a new association is made. Identifier associations are timestamped, making it possible to see the naming history of any sequences.

For new sequences, SeqRepo generates a set of computed identifiers using computational digests and loads these as with conventional identifiers. Currently, these digests are sha512t24u, MD5, SEGUID, and SHA-1.

### Snapshots

Snapshots are created using hard links in the file system. Therefore, new snapshots consume only the space required for the incremental BGZF and database files. Similarly, transfers with rsync transfer only the incremental changes.

### Interfaces

SeqRepo includes two interfaces: the native Python interface, and a read-only REST-based interface to be used for local access by programs written in other languages. The SeqRepo package also implements the GA4GH refget protocol to support clients that require that interface. The refget implementation passes the compliance suite provided by the authors. Table 2 summarizes operations supported by these three interfaces.

**Table 1.**
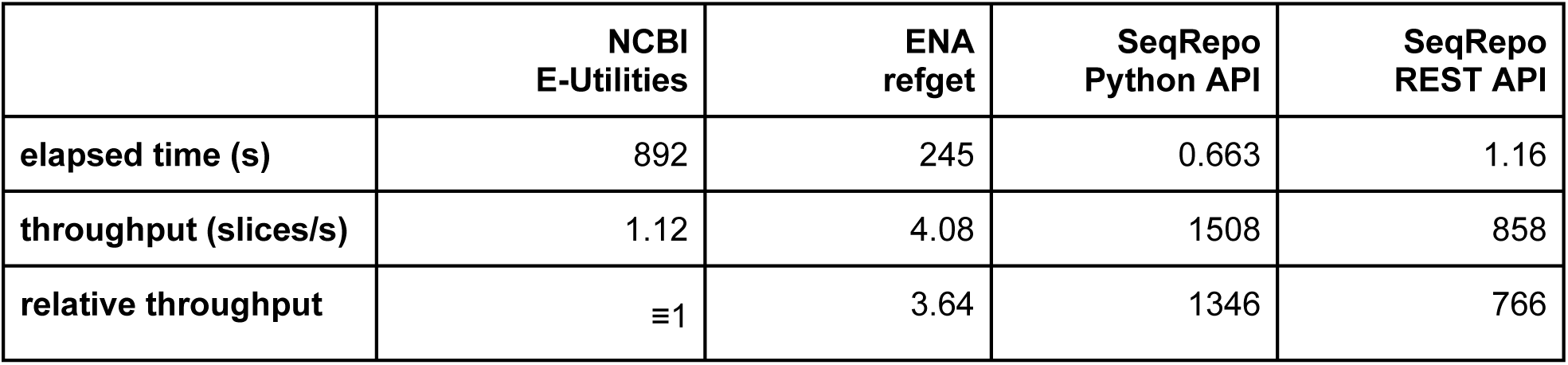
Timing results for remote and local sequence sources. Timings are for 1,000 sequence lookups from 1) NCBI nucleotide sequences using the E-utilities interface, 2) European Nucleotide Archive using the refget protocol, 3) Local SeqRepo using the native Python interface, 4) Local SeqRepo interface using a SeqRepo REST API. Local SeqRepo access offers the best performance; using the SeqRepo REST API adds overhead, but enables access from other programming languages. Details are provided in S3 Code.

|  | NCBI<br>E-Utilities | ENA<br>refget | SeqRepo<br>Python API | SeqRepo<br>REST API |
| --- | --- | --- | --- | --- |
| elapsed time (s) | 892 | 245 | 0.663 | 1.16 |
| throughput (slices/s) | 1.12 | 4.08 | 1508 | 858 |
| relative throughput | ≡1 | 3.64 | 1346 | 766 |

**Table 2.**
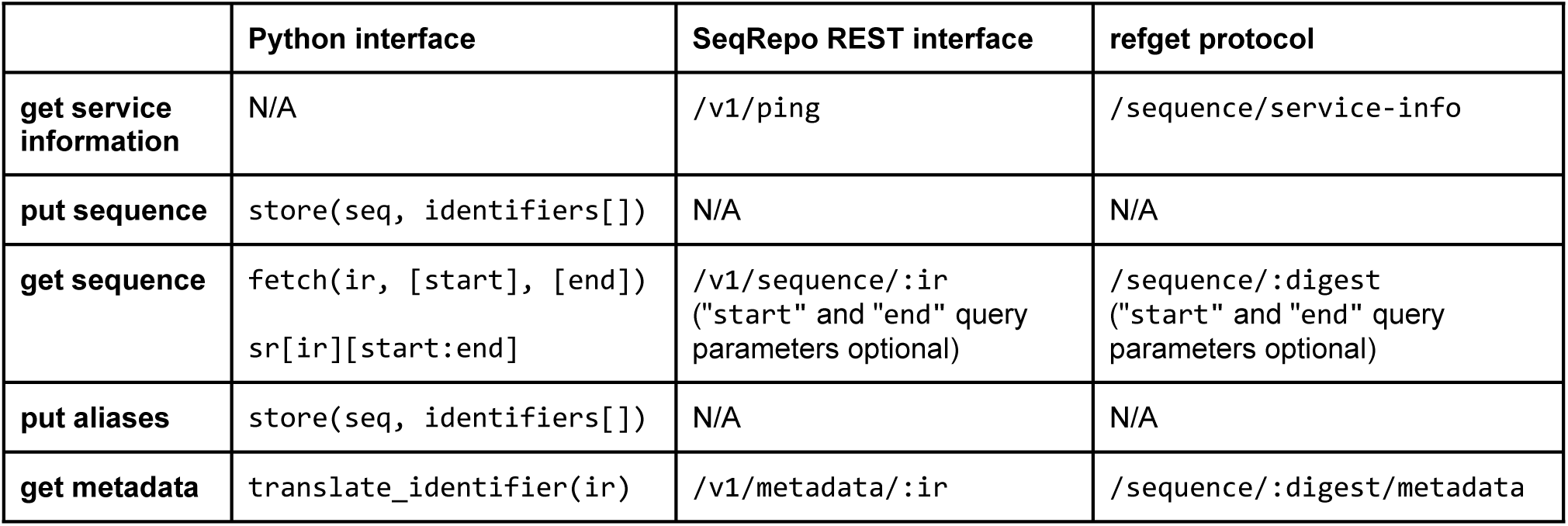
Summary of operations provided by the Python interface, REST interface, and refget protocol interface. All interfaces support rapid access to slices of chromosome-sized sequences. The SeqRepo provides two mechanisms to fetch sequence slices: a fetch() method, and a dict-style access that permits a SeqRepo instance to be accessed as a Python dictionary. The SeqRepo REST interface and refget protocol are read-only interfaces. “ir” denotes an identifier of the form *namespace:alias*; an alias may be used without namespace if it is globally unique. The refget protocol itself currently requires the use of digests for queries.

SeqRepo interfaces represent identifiers as W3C Compact URIs (CURIEs), mapping *<namespace,alias>* to the *<prefix, reference>* nomenclature; the converse operation is also supported when CURIE-formatted identifiers are provided as sequence identifiers.

### Installation Scenarios

SeqRepo supports four installation scenarios:

1. A read-write instance, loaded and maintained locally with the seqrepo command line interface.
2. A read-only instance of the human collection, mirrored and updated using the seqrepo command line tool.
3. A docker data-only container for linking with other application containers. This approach is useful to share a single data container with multiple docker applications.
4. SeqRepo and refget REST interfaces, also available as a docker image.

Detailed instructions for each of these options are available at the GitHub repository (https://github.com/biocommons/biocommons.seqrepo).

## Results

### sha512t24 Timing and Statistical Analysis

Members of the SHA-1, SHA-2, and MD5 family of digests were compared for timing. Because SHA-512 is specified in terms of 64-bit operations, the increased complexity of this algorithm is offset by the ability to process data twice as fast as with 32-bit operations specified by shorter digest algorithms. SHA-512 provides 512 bits of digest, which is far more than required for even extremely large sets of messages. Because this digest will be used as a key and transmitted widely, a practical tradeoff between key size and collision probability is desirable. Base64 results in encodings that are ceil(4/3) of the size of the input message; therefore, the truncation length should be modulo 3 for efficiency. The probability of collision was evaluated for a series of modulo-3 truncation lengths, and 24 bytes was chosen as a compromise between number of expected messages (sequences) and collision probability. For example, in a corpus of 10e+18 sequences, a key size of 24 bytes is expected to result in a collision probability of <1e-21. See S1 Code for details on the timing and statistical analyses. Note that the truncated digest used here is not the same as the truncated digest proposed in the SHA-2 family of algorithms; in particular, we use the SHA-512 initialization vector in order to enable prefix abbreviations for larger digests.

### Human SeqRepo Collections

SeqRepo may be used for sequences of any type from any species. The current Human releases (available on https://dl.biocmmons.org/) consume approximately 12 GB on disk (with indexes) and consist of over 925,000 unique sequences from NCBI35, NCBI36, GRCh37 (with patches), GRCh38 (with patches), JRGv1, JRGv2, RefSeq (NM, NP, XM, XP, NG), and Ensembl (ENST, ENSP). The differential size between snapshots is approximately 1 GB, and is easily incrementally updated using the seqrepo command line interface. The command line interface enables users to easily import fasta files into custom sequence collections.

### SeqRepo Python interface

SeqRepo provides a facile interface for accessing sequences, demonstrated in Figure 2.

**Figure 2.**
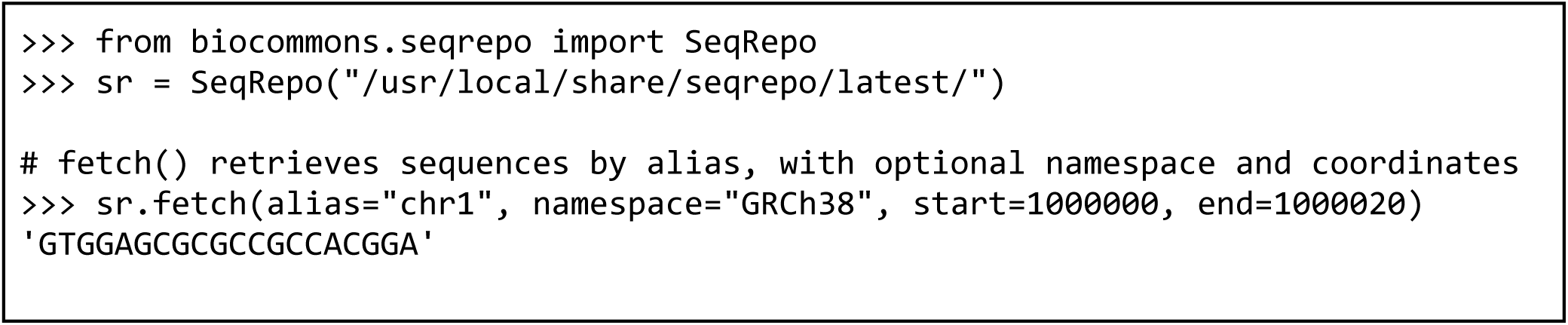
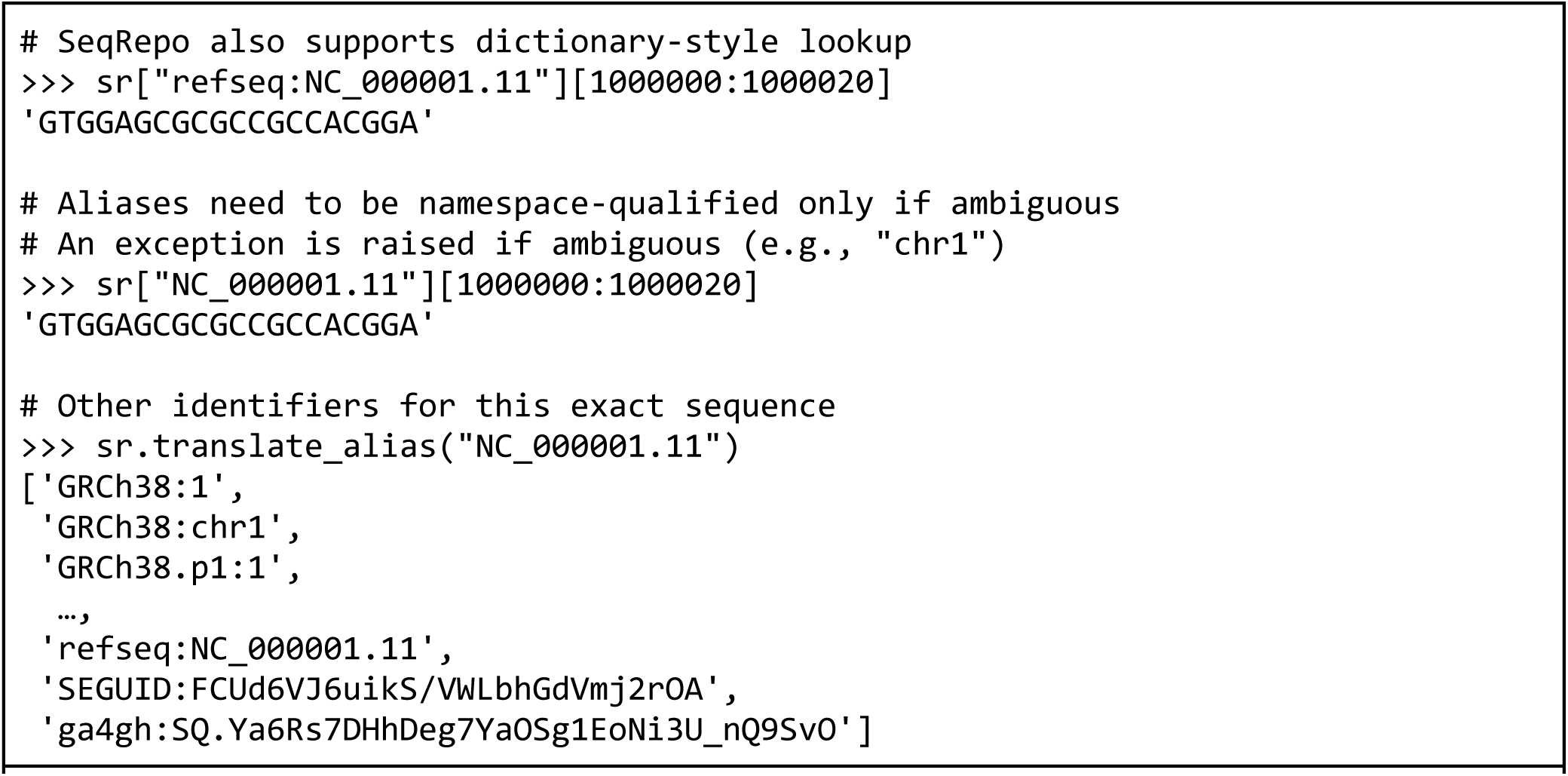
Examples of the native Python interface. SeqRepo retrieves sequences and metadata using conventional identifiers (*i*.*e*., from NCBI, Ensembl, GRCh, LRG, and other sources) and from digest identifiers (*i*.*e*., sha512t24u, ga4gh, md5, SEGUID). identifiers are namespaced, and generally written as “<namespace>:<alias>“. Aliases are always unique within a namespace. See the SeqRepo repository for installation instructions and S2 Code for example details.

### SeqRepo REST API

SeqRepo also provides a REST API to access sequence slices and metadata from any language while preserving the performance advantages of local sequence collections. An example of using the REST interface from a Linux command line is shown in Figure 3.

**Figure 3.**
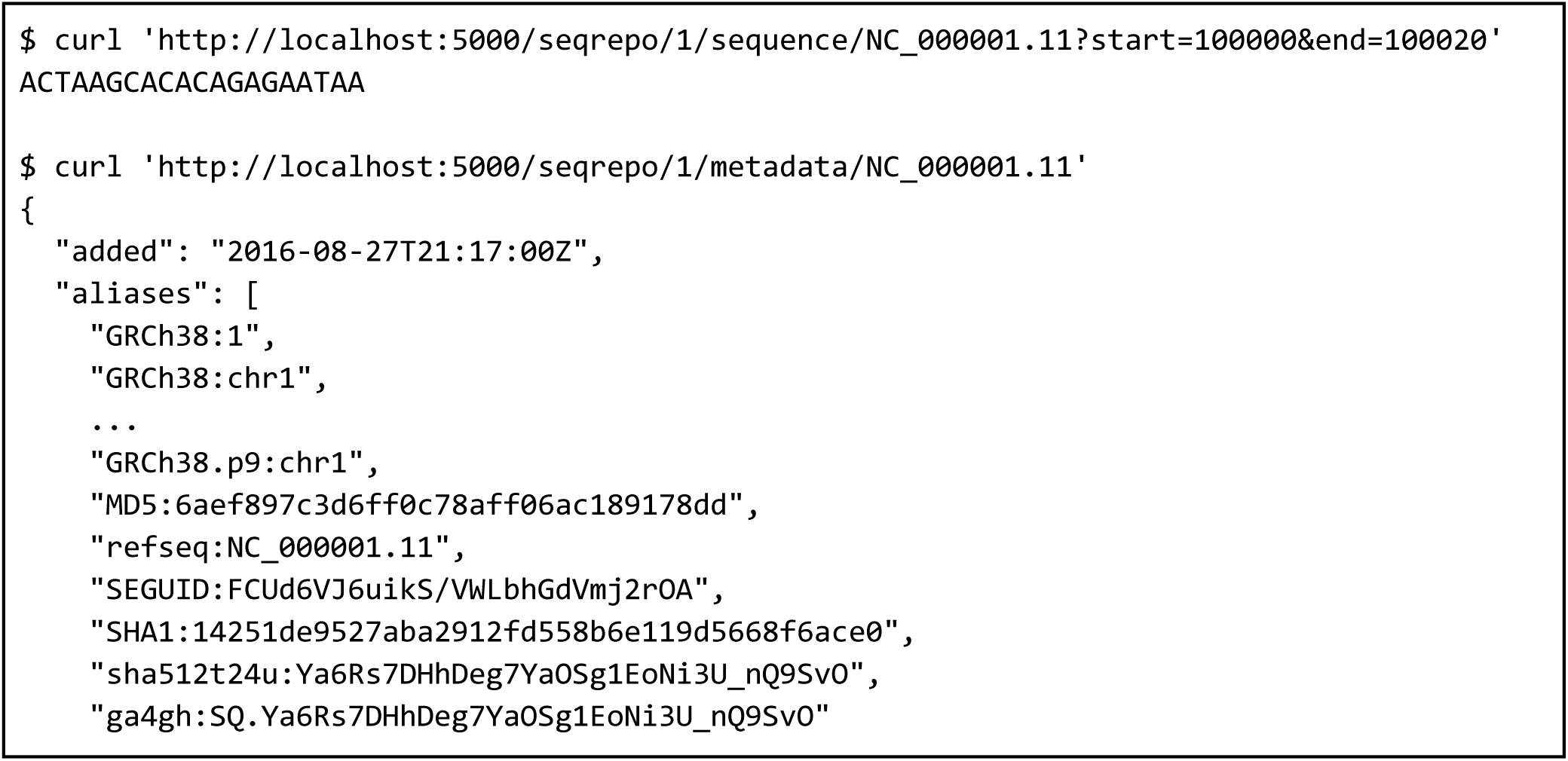
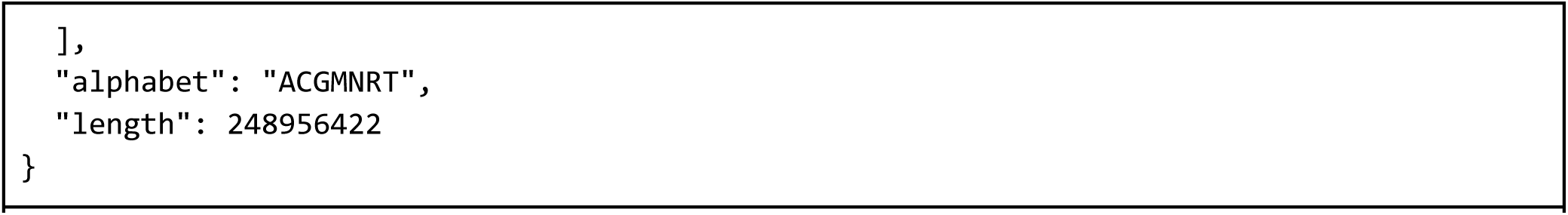
Examples of the SeqRepo REST API. See the SeqRepo repository for installation instructions and S2 Code for example details.

### Timing Comparison with NCBI E-Utilities and ENA refget

In order to demonstrate the impact of local sequence storage on processing speed, we measured elapsed time for fetching 1,000 random sequence slices of 1-30 nucleotides from all chromosomes of GRCh38 using NCBI E-utilities (Sayers, 2010), the European Nucleotide Archive implementation of refget, and SeqRepo. Results are showing in Table 1.

SeqRepo tests were performed on a modern laptop with a solid-state disk. NCBI E-utilities and ENA refget tests were performed on a AWS c4.large instance in us-east.

NCBI imposes rate limiting of 3 queries per second without an API key, or 10 queries per second with an API key, which we used. It was also necessary to implement rate limiting and retry logic in order to successfully retrieve all sequences. A genomic slice can be retrieved in one query, resulting in a theoretical maximum throughput of 10 sequence slices/second. Server-side rate limiting also defeats client-side parallelism.

Given the magnitude of the differences between these methods, we did not seek higher precision timings. See Supplementary Information for methods and timing data.

## Conclusion

SeqRepo permits the use of conventional identifiers and digests for accessing and retrieving sequences. It is not necessary to adopt digest-based identifiers when using SeqRepo. This difference makes it possible to easily translate between conventional identifiers and digest identifiers.

Furthermore, a locally-maintained SeqRepo instance enables pipelines to transparently mix public and custom sequences, such as masked sequences or alternative assemblies for variant calling.

Since its release in 2016, we have released 19 snapshots and improved namespace support for the GA4GH Variation Representation Specification. While refget and SeqRep can translate from GA4GH sequence identifiers to conventional aliases, SeqRepo is currently the only service that can translate from conventional identifiers to GA4GH sequence identifiers. This service is essential for translating variation to GA4GH standards.

Circular sequences are not directly supported by SeqRepo. However, such sequences may be stored in a linear form. We anticipate adding support for circular sequences in the future. SeqRepo currently implements only the BGZF backend described here. However, generalizing the interface to REDIS, AWS Aurora, or AWS S3 (as used by refget) would be straightforward and is under consideration.

Deciding between remote and local sequence services requires considerations of runtime performance, maintenance effort, data currency requirements, data privacy, and overall system availability. We demonstrated a 600-fold performance improvement for using local sequence storage rather than remote services. Although the performance benefit of using local sequences is unsurprising, these magnitude of the difference provides guidance regarding this important choice when designing an analysis pipeline, particularly if high-throughput is desired. SeqRepo significantly lessens the effort required to maintain a local sequence repository and provides significant performance benefits over remote sequence lookup services.

## Acknowledgements

SeqRepo was initially funded by Invitae.

## Supporting Information

- S1 Code. <u>SHA-512 timing and collision analysis (jupyter notebook)</u>
- S2 Code. <u>API Examples (jupyter notebook)</u>
- S3 Code. <u>Timing Comparisons (jupyter notebook)</u>

